# Ultrasensitive Detection of Alpha-Synuclein Oligomers in Human Plasma Using Optimized Nano-QuIC

**DOI:** 10.64898/2026.02.04.703848

**Authors:** Hyeonjeong Jeong, Peter R. Christenson, Hyerim Ahn, Hilal A. Lashuel, Peter A. Larsen, Sang-Hyun Oh, Hye Yoon Park

**Affiliations:** Department of Electrical and Computer Engineering, University of Minnesota, Minneapolis, Minnesota 55455, United States; Laboratory of Molecular and Chemical Biology of Neurodegeneration, Brain Mind Institute, Ecole Polytechnique Federale de Lausanne (EPFL), Lausanne, Switzerland; ND BioSciences SA, Switzerland; Weill Cornell Medicine Qatar, Education City, Qatar Foundation, Doha, Qatar; Minnesota Center for Prion Research and Outreach, University of Minnesota, St. Paul, Minnesota 55108, United States; Department of Veterinary and Biomedical Sciences, University of Minnesota, St. Paul, Minnesota 55108, United States

## Abstract

Early diagnosis of Parkinson’s disease (PD) is critical, as clinical symptoms typically emerge only after substantial neuronal loss. While α-synuclein (α-Syn) oligomers in blood are promising biomarkers for early detection, their clinical utility is limited by their low abundance and the presence of inhibitory components in the plasma matrix. To address these limitations, we tailored the Nanoparticle-enhanced Quaking-Induced Conversion (Nano-QuIC) platform specifically for the ultrasensitive detection of α-Syn oligomers in human plasma. We identified critical reaction determinants by investigating buffer pH, ionic strength, detergent types, and shaking conditions. Furthermore, the integration of silica nanoparticles (siNPs) proved essential in mitigating plasma matrix interference, ensuring robust and reproducible protein aggregation. Under these optimized conditions, the assay achieved a detection limit of 100 pg/mL for α-Syn oligomers spiked into human plasma. These results demonstrate that our adapted Nano-QuIC platform provides a highly sensitive and minimally invasive method for detecting pathological α-Syn species, offering a significant advancement toward the development of early-stage PD diagnostics.

## Introduction

Parkinson’s Disease (PD) is a neurodegenerative disease that causes motor dysfunction and cognitive impairment. The pathological hallmark of PD is the misfolding and aggregation of α-synuclein (α-Syn), a presynaptic protein that accumulates as oligomeric and fibrillar aggregates, forming the Lewy bodies (LBs) and Lewy neurites (LNs). Mounting evidence points to the formation of LBs and LNs as key contributors to the progressive loss of dopaminergic neurons in PD, although the nature of the primary pathogenic species remains a subject of active research and debate (1). Research suggests that α-Syn misfolding and aggregation begin up to eight years before the clinical manifestation of PD (2), underscoring the need for early diagnostic approaches to enable timely therapeutic intervention (1,3).

Recent advances in seed amplification assays (SAAs) have significantly enhanced the detection of misfolded α-Syn fibrils, with some studies demonstrating the ability to detect concentrations as low as 4 pg/mL using preformed fibrils (PFF) in buffer as reference seeds (4). Despite these successes, the application of these assays for early-stage diagnosis is often hampered by a lack of reliable biomarkers that appear at the very onset of the disease. In this context, α-Syn oligomers have emerged as highly promising early-stage biomarkers (3,5–7). Oligomers are believed to appear early in the aggregation cascade, preceding the formation of mature fibrils and LBs. (Fig. 1a) (8,9). Their pathological role is linked to several types of cellular malfunction, including membrane disruption, mitochondrial impairment, synaptic dysfunction, and neuroinflammation (10).

**Figure 1.**
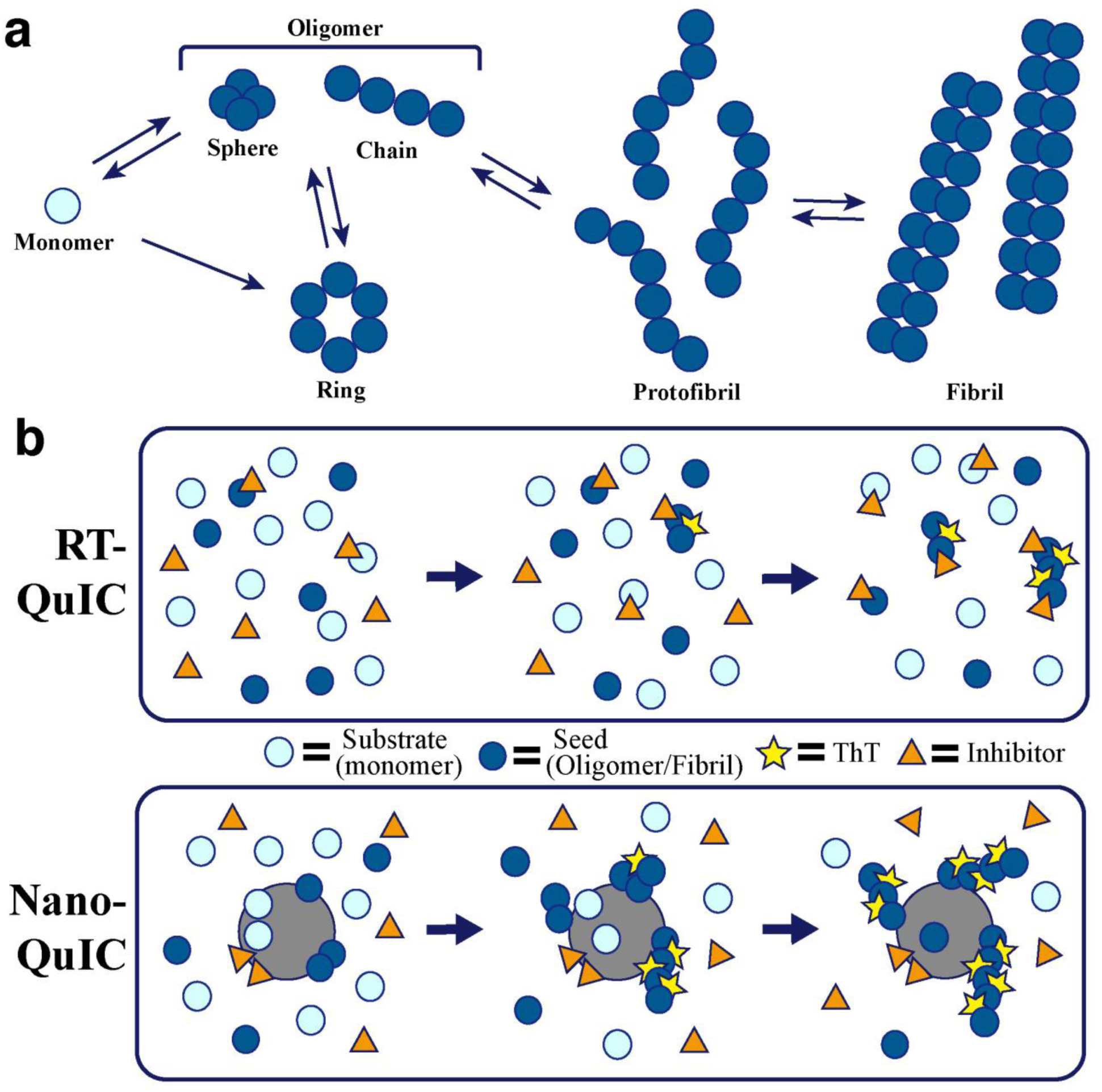
Overview of misfolding and QuIC assay for oligomer detection. **(a)** α-Syn aggregation pathway showing the transition from monomers to various oligomeric forms (spheres, rings, chains), which can further convert into protofibrils and mature fibrils. **(b)** Comparison of conventional RT-QuIC and Nan-QuIC assays. In RT-QuIC, the presence of inhibitors in plasma can interfere with the interaction between monomeric substrates and aggregation seeds (oligomers/fibrils), reducing assay efficiency. In contrast, Nano-QuIC incorporates silica nanoparticles, which help concentrate seeds and substrates while mitigating the effects of inhibitors, thereby enhancing aggregation kinetics and ThT fluorescence signal detection.

Although their precise role in the aggregation pathway remains under investigation, numerous studies support a model in which α-Syn monomers transitions through structurally heterogeneous oligomers before forming protofibrils and fibrils (Fig. 1a) (8,11,12). The structural importance of oligomers has gained further attention following recent studies demonstrating abundant oligomer accumulation in the brains of LB-negative *LRRK2* mutation carriers (13,14). These findings provide a possible explanation for how individuals may develop classic PD symptoms even in the absence of traditional Lewy pathology. While α-Syn oligomers have been detected at elevated levels in both the brains and cerebrospinal fluid (CSF) of patients with various synucleinopathies, including PD and dementia with Lewy bodies (DLB) (10), CSF-based diagnostics are limited by the invasive nature of lumbar punctures. This invasiveness makes CSF sampling unsuitable for routine population screening or frequent longitudinal monitoring.

Consequently, there is a strong shift toward utilizing peripheral fluids such as blood, saliva, and urine, which are more accessible and patient-friendly (5,6,15–21). Blood is particularly attractive due to its widespread clinical use and potential for longitudinal sampling. However, blood-based α-Syn detection faces significant technical barriers. The majority of α-Syn in blood originates from red blood cells (RBCs), and hemolysis during sample collection or processing can drastically skew α-Syn levels in plasma or serum (6,22). Furthermore, blood contains numerous inhibitory components that interfere with the efficiency of SAAs, particularly when targeting low-abundance species such as oligomers (23).

To adapt SAAs, which has shown promise in CSF and some peripheral tissues (24–32), to this complex blood matrix, various strategies have been explored. For instance, immunoprecipitation-based RT-QuIC (IP/RT-QuIC) (19) and other blood-based SAAs (33,34) have yielded promising diagnostic results in both prodromal and symptomatic stages. However, these methods often remain labor-intensive or limited by plasma matrix interference. To address these challenges, Nano-QuIC was recently developed to mitigate inhibition and accelerate nucleation using silica nanoparticles (siNPs) (Fig. 1b) (35). Yet, while Nano-QuIC has demonstrated superior speed and sensitivity for detecting α-Syn fibrils, its capacity to detect oligomeric seeds has not yet been systematically evaluated.

In this study, we sought to bridge this gap by optimizing the Nano-QuIC assay for the ultrasensitive detection of α-Syn oligomers in human plasma. We systematically evaluated the impact of critical assay parameters, including buffer pH, ionic strength (Na⁺ concentration), detergent type, and shaking protocols, on the seeding activity of α-Syn oligomers. All α-Syn species (monomers, oligomers, and fibrils) utilized in this study were thoroughly characterized for their structural properties and stability. Our results confirm that the oligomer seeds maintain their structural integrity under the optimized conditions, ensuring that the observed assay performance is driven by stable oligomeric species rather than pre-existing or spontaneously formed fibrils. Ultimately, this work establishes a robust assay platform capable of ultrasensitive detection of α-Syn oligomers, providing a vital foundation for the development of minimally invasive diagnostics for early-stage PD.

## Results

### Comparison of RT-QuIC and Nano-QuIC for misfolded α-Syn detection

The addition of siNPs to α-Syn seed amplification assays has previously been shown to accelerate reaction kinetics by adsorbing and concentrating α-Syn substrate on their surface, and reducing the inhibitory effects of components present in human plasma (35). To validate these findings in our system, we performed side-by-side comparisons of standard RT-QuIC and Nano-QuIC assays (Fig. 1b). To start, all experiments were conducted in three replicates under identical reaction conditions at pH 7.4, 339 mM Na⁺, 0.002% sodium dodecyl sulfate (SDS), and a 60 s shake / 60 s rest cycle at 42°C. All α-Syn species (monomers, oligomers, and fibrils) used in this study were prepared and extensively characterized by ND Biosciences (Supplementary Datasheet S1). We independently verified that the oligomer sizes ranged from 14 to 33 nm (Fig. S1a), and the PFF lengths were between 51 and 362 nm (Fig. S1b), consistent with the manufacturer’s specifications.

As shown in Fig. 2, Thioflavin T (ThT) fluorescence was continuously monitored in reactions seeded with either 9 μg/mL of α-Syn oligomers or 90 μg/mL of PFFs spiked into human plasma. To normalize the concentration of active seeding units rather than total protein mass, we used a lower mass of oligomers, as their smaller sizes result in a higher number of seeds per unit mass compared to the longer PFFs. To quantitatively compare assay performance, we calculated two key metrics: the rate of amyloid formation (RAF), defined as the inverse of the time required to reach a predefined fluorescence threshold, and the maximum point ratio (MPR), defined as the ratio of the peak fluorescence intensity to the baseline (36).

**Figure 2.**
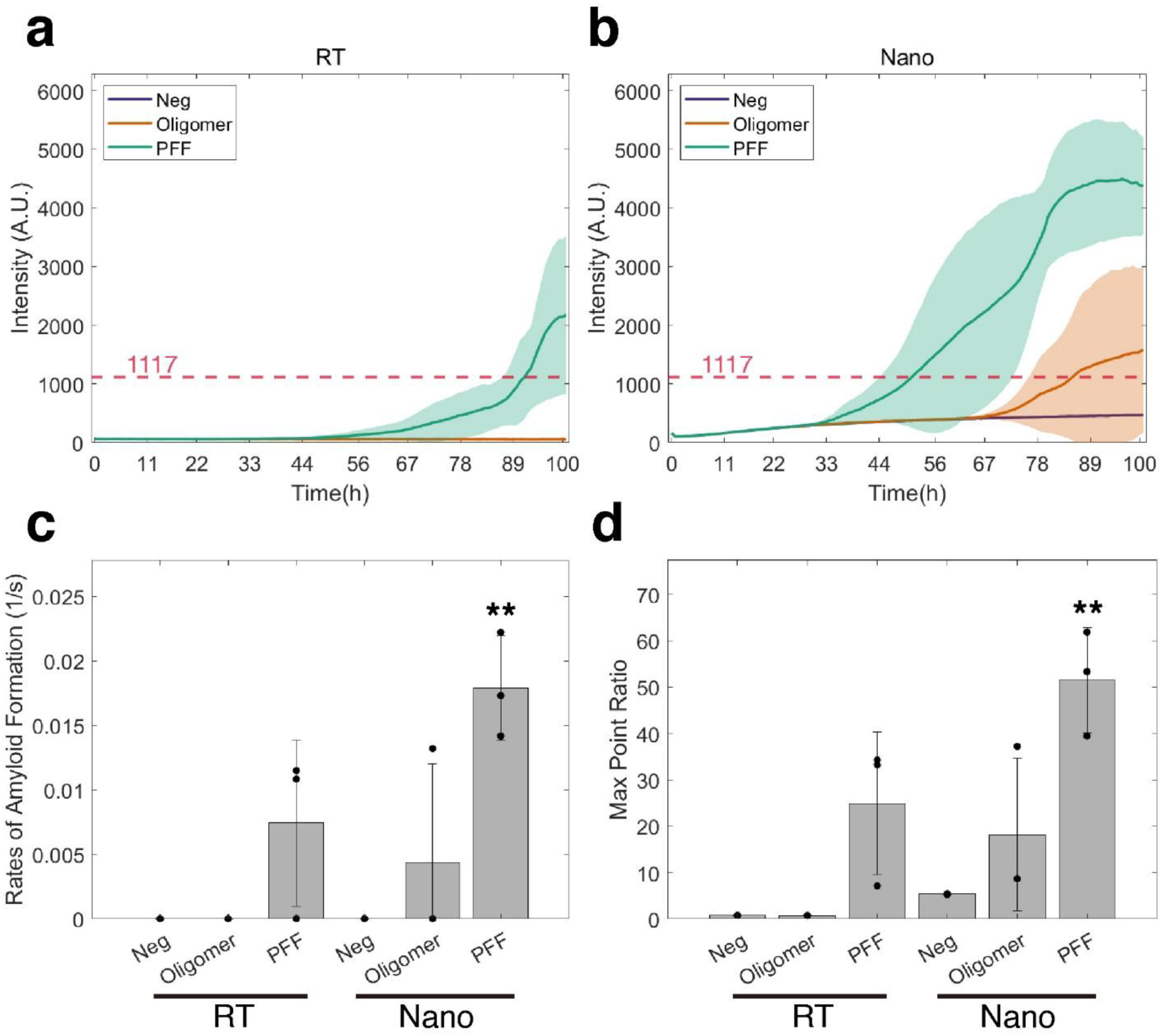
RT-QuIC vs Nano-QuIC for α-Syn oligomer and PFF detection. Average ThT fluorescence curves of **(a)** RT-QuIC and **(b)** Nano-QuIC assay. Solid lines represent the mean, and shaded areas indicate ± SD from three replicates of plasma samples seeded with 9 μg/mL of α-syn oligomer (orange) and 90 μg/mL of PFF seeds (green). The red dashed line indicates the threshold for positive detection. **(c)** RAF and **(d)** MPR of negative controls and seeded plasma samples. Data points represent individual replicates (n = 3), and bars indicate mean ± SD. Statistical comparisons were made between the negative controls and other cases under the same experimental conditions. Statistical significance is denoted as *p < 0.05, **p < 0.01, and ***p < 0.001.

As illustrated in Fig. 2a and 2b, the Nano-QuIC assay exhibited substantially faster fibril amplification kinetics compared to the conventional RT-QuIC. When seeded with PFFs, Nano-QuIC initiated fibril amplification at approximately 30 hours, whereas RT-QuIC showed a detectable signal only after 50 hours. In oligomer-seeded reactions, while RT-QuIC remained completely unresponsive throughout the assay period, Nano-QuIC successfully captured the seeding activity in a subset of replicates (1/3) starting around 70 hours. Notably, no spontaneous fibril formation was observed in the monomer-only conditions (Neg) for either platform. Quantitative analysis further supported these kinetic advantages. As illustrated in Fig. 2c, Nano-QuIC yielded significantly higher RAF values in PFF-seeded reactions and demonstrated a unique capacity for detecting oligomeric seeds, unlike RT-QuIC. Furthermore, MPR values were markedly elevated in Nano-QuIC for both PFF and oligomer seeds (Fig. 2d), indicating stronger fluorescence signals and improved assay sensitivity.

Our results demonstrate that the Nano-QuIC assay exhibits enhanced sensitivity and greater resistance to the inhibitory components in human plasma compared to the standard RT-QuIC. This underscores the potential of the Nano-QuIC platform for the robust detection of α-Syn seeds in clinical matrices and establishes a strong foundation for further assay optimization.

### Optimized buffer pH for misfolded α-Syn oligomer detection

To determine the optimal buffer conditions for α-Syn oligomer and PFF seeds, we evaluated the performance of the Nano-QuIC assay using various buffers across a range of pH values at 339 mM Na⁺ concentration, 0.002% SDS, and a 60 s shake / 60 s rest cycle at 42°C. pH is known to be one of the key factors influencing α-Syn aggregation, with acidic conditions generally accelerating aggregation (37,38). We used different buffers to achieve a range of pH conditions: phosphate buffer at pH 5.8, Tris buffer at pH 6.5, phosphate-buffered saline (PBS) at pH 7.4, and phosphate buffer at pH 8.2. The experiment was conducted with three replicates. Fig. 3a-d present the ThT fluorescence kinetics of Nano-QuIC reactions with 9 μg/mL of α-Syn oligomers and 90 μg/mL PFF seeds spiked into plasma, each conducted in buffers with the indicated pH values. As the buffer pH increased, the fluorescence amplitude decreased, likely reflecting the pH sensitivity of ThT, whose fluorescence is enhanced under more acidic conditions (39). For oligomer seeds, fibril formation began at approximately 10 hours at pH 5.8 and 6.5 hours at pH 6.5, whereas at pH 7.4, amplification was delayed until around 70 hours (Fig. 3a-c). No detectable fibril amplification was observed at pH 8.2 (Fig. 3d). Similarly, fibril amplification of α-Syn PFF seeds began at ∼7 hours at pH 5.8 and ∼5.5 hours at pH 6.5, was delayed to ∼33 hours at pH 7.4, and was absent at pH 8.2 (Fig. 3a-d). Consistent with these trends, RAF values were higher at pH 5.8 and pH 6.5, indicating faster aggregation kinetics under acidic conditions (Fig. 3e).

**Figure 3.**
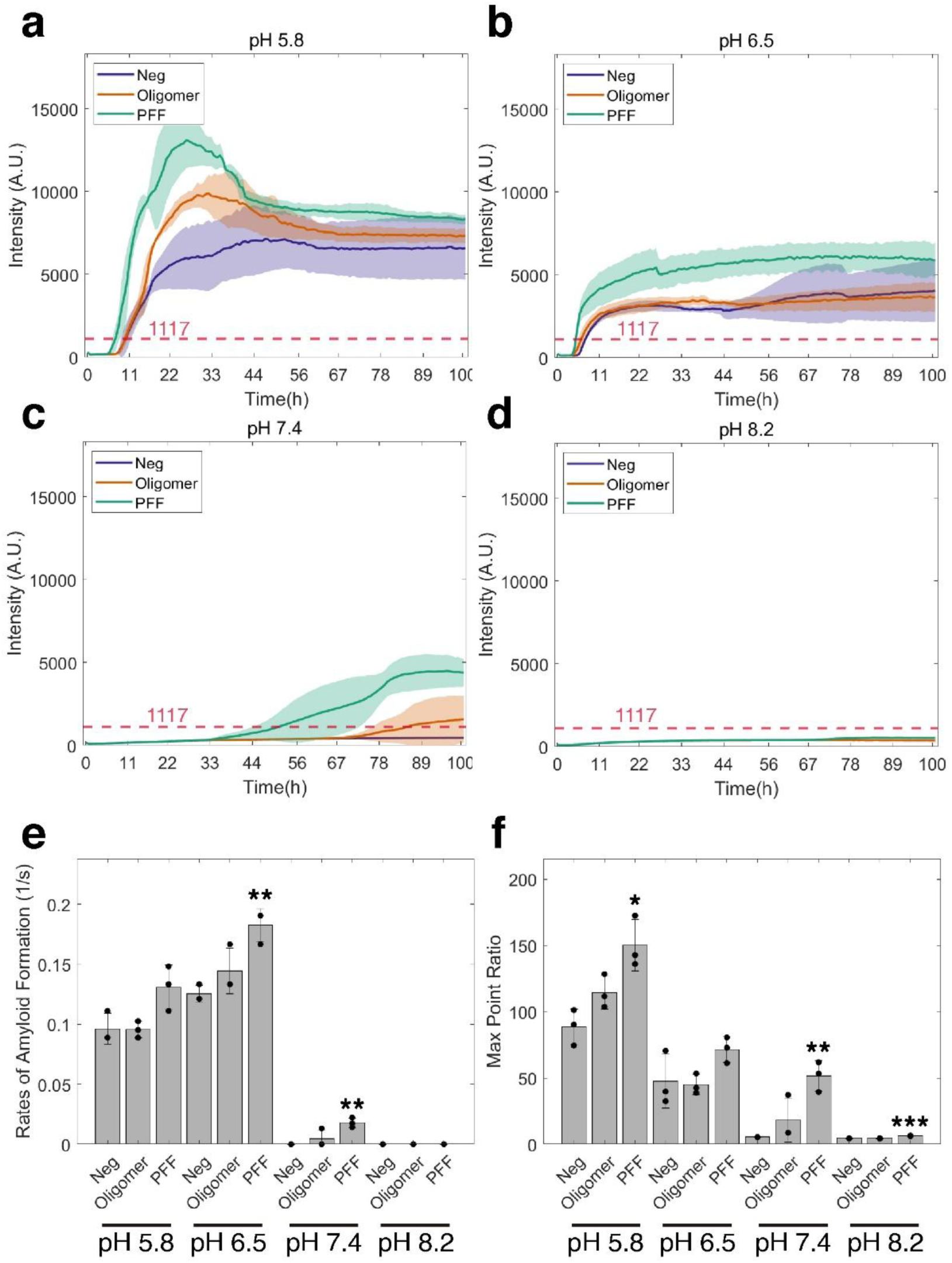
Effect of pH on the detection of α-Syn oligomer and PFF seeds. Average ThT fluorescence curves of Nano-QuIC assay on **(a)** pH 5.8, **(b)** 6.5, **(c)** 7.4, and **(d)** 8.2. Solid lines represent the mean, and shaded areas indicate ± SD from three replicates of plasma samples seeded with 9 μg/mL of α-syn oligomer (orange) and 90 μg/mL of PFF seeds (green). The red dashed line indicates the threshold for positive detection. **(e)** RAF and **(f)** MPR of negative controls and seeded plasma samples. Data points represent individual replicates (n = 3), and bars indicate mean ± SD. Statistical comparisons were made between the negative controls and other cases under the same experimental conditions. Statistical significance is denoted as *p < 0.05, **p < 0.01, and ***p < 0.001.

However, these low pH conditions promoted spontaneous aggregation in monomer-only samples (Neg), which compromised assay specificity. At pH 5.8, no significant difference was observed between the monomer-only condition and any seeded reactions. At pH 6.5 and pH 7.4, a significant difference was observed between PFF-seeded reactions and the monomer-only condition, whereas oligomer-seeded reactions were not significantly different. No signal was detected at pH 8.2.

In Fig. 3f, MPR values were also highest at lower pH. However, spontaneous aggregation in the monomer-only condition limited the interpretability of these high MPR values. Although MPR values were higher at pH 5.8 and pH 6.5, the dominance of spontaneous aggregation reduced discrimination between seeded and unseeded samples. At pH 5.8, only a small significant difference was observed between the PFF-seeded samples and the monomer-only condition, and at pH 6.5, no significant differences were observed. In contrast, pH 7.4 produced slightly lower MPR values but showed minimal spontaneous aggregation in the monomer-only condition, enabling clear discrimination between PFF-seeded samples and monomer-only samples. While the difference for oligomer-seeded samples did not reach statistical significance, they remained visually distinguishable from the monomer-only condition, suggesting improved specificity at pH 7.4.

Overall, PBS at pH 7.4 yielded the most robust and consistent ThT fluorescence signals across replicates for PFF seeds. Although oligomer-seeded reactions did not reach statistical significance under these conditions, the observed trends were highly promising. These findings indicate that pH 7.4 provides an optimal balance between aggregation sensitivity and assay specificity and was therefore selected as the primary buffer condition for subsequent Nano-QuIC assays.

### Optimized Na^+^ concentration for misfolded α-Syn oligomer detection

Next, we optimized the Na⁺ concentration in the Nano-QuIC assay, as Na⁺ is known to affect α-Syn aggregation by modulating electrostatic interactions involved in fibril formation (40). The total Na⁺ concentration, including contributions from the PBS buffer, was calculated to ensure accurate control of ionic strength in the assay. We tested four final Na⁺ concentrations, 169.33 mM, 300 mM, 500 mM, and 700 mM, using an assay performed at pH 7.4 with 0.002% SDS and a 60 s shake / 60 s rest cycle at 42°C.

Starting with this experiment, we incorporated a pre-analytical centrifugation step to mitigate the effects of plasma inhibitors and improve sensitivity (35). Plasma samples spiked with 100 ng/mL of α-Syn oligomers or 1 μg/mL of PFFs were centrifuged at 21,000 × g for 40 min at 4°C. The supernatant was discarded to remove soluble inhibitors, and the pellet was resuspended in 14 µL of detergent. This enrichment process effectively concentrated the seeds, allowing us to use initial spike concentrations approximately 90-fold lower than those used in the previous experiments.

Fig. 4 shows the ThT fluorescence kinetics of Nano-QuIC reactions for both oligomer- and PFF-spiked plasma across various Na⁺ concentrations. To evaluate assay performance, we monitored the average fluorescence intensity and determined the RAF and MPR across three replicates. As shown in Fig. 4a-d, increasing Na⁺ concentration led to an enhanced ThT fluorescence intensity in PFF-seeded reactions. For oligomer-seeded reactions, a notable increase in aggregation was observed at 500 mM, with high consistency across replicates. Quantitative analysis in Fig. 4e reveals that RAF values for both oligomers and PFFs increased at higher Na⁺ concentrations, indicating accelerated aggregation kinetics. Notably, PFF-seeded reactions showed statistically significant differences from negative controls at most concentrations, except for 300 mM. In contrast, for oligomer seeds, only the 500 mM Na⁺ concentration showed a significant difference from the negative controls. Regarding the MPR (Fig. 4f), a significant increase was observed in PFF-seeded reactions at 700 mM Na⁺, whereas no significant differences were found for oligomer seeds at any tested concentration. At 500 mM Na⁺, only one out of three negative control replicates exhibited a spontaneous signal, suggesting this concentration provides an optimal balance for assay sensitivity and specificity. Consequently, 500 mM was selected as the optimal Na⁺ concentration for oligomer detection in the Nano-QuIC assay.

**Figure 4.**
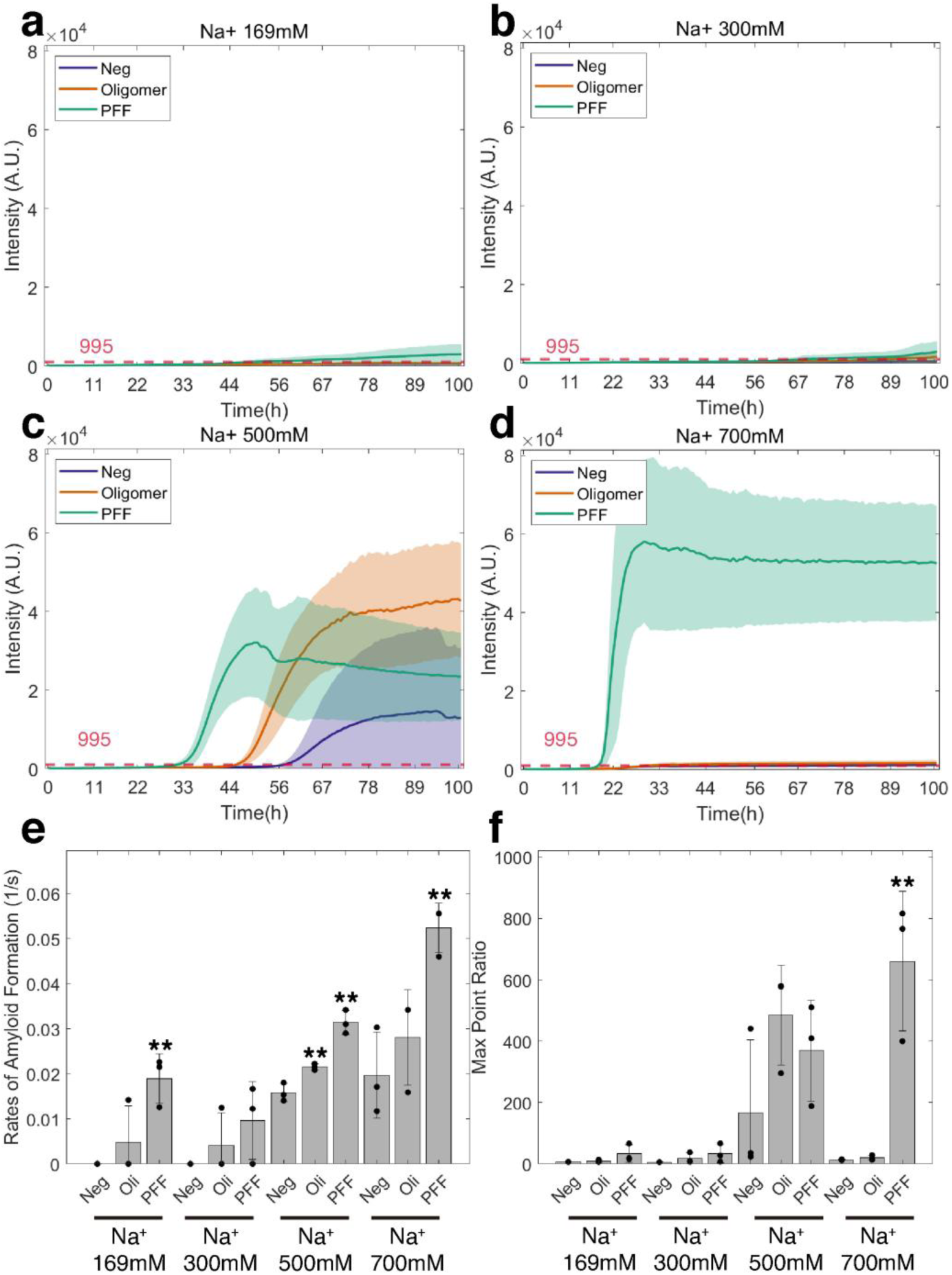
Effect of Na^+^ concentration on the detection of oligomer and PFF seeds. Average ThT fluorescence curves of Nano-QuIC assay with **(a)** 169 mM, **(b)** 300 mM, **(c)** 500mM, and **(d)** 700 mM Na^+^ concentrations. Solid lines represent the mean, and shaded areas indicate ± SD from three replicates of plasma samples seeded with 100 ng/mL of α-syn oligomer (orange) and 1 μg/mL of PFF seeds (green). The red dashed line indicates the threshold for positive detection. **(e)** RAF and **(f)** MPR of negative controls and seeded plasma samples. Data points represent individual replicates (n = 3), and bars indicate mean ± SD. Statistical comparisons were made between the negative controls and other cases under the same experimental conditions. Statistical significance is denoted as *p < 0.05, **p < 0.01, and ***p < 0.001.

### Optimized detergent type for misfolded α-Syn oligomer detection

We then compared two detergents, SDS and Triton X-100, to evaluate their effects on the seeding efficiency of α-Syn oligomers and PFFs in the Nano-QuIC assay. Based on previous optimization, all reactions were performed with 500 mM Na⁺ at pH 7.4, using a 60 s shake / 60 s rest cycle at 42°C (n = 4). We utilized the same enrichment protocol as described above for plasma samples seeded with 100 ng/mL of α-Syn oligomers or 1 μg/mL of PFFs. The final detergent concentrations were 0.002% for SDS and 0.02% for Triton X-100. SDS is a commonly used detergent in SAAs, known to accelerate α-Syn aggregation by modulating electrostatic and hydrophobic interactions (41). Triton X-100 is another detergent that has recently been reported to accelerate the RT-QuIC assay for detecting misfolded α-Syn across diverse clinical samples (42).

As shown in Fig. 5a and 5b, both SDS and Triton X-100 enabled the detection α-Syn oligomer-and PFF-mediated seeding activity. However, oligomer-seeded reactions exhibited significantly stronger fibril formation with Triton X-100, as shown in Fig. 5b. α-Syn oligomers-mediated seeding and fibrillization typically began at approximately 69 hours with SDS and around 40 hours with Triton X-100. For PFF-seeded reactions, fibril amplification started at approximately 50 hours with SDS and 36 hours with Triton X-100. Although Triton X-100 showed a tendency for slightly increased spontaneous nucleation compared to SDS, the negative controls remained clearly distinguishable from both oligomer- and PFF-seeded reactions (Fig. 5c). Conversely, SDS-based reactions exhibited high variability, making it difficult to consistently distinguish between conditions (Fig. 5a and 5d). Quantitatively, Triton X-100 yielded significantly higher RAF and MPR values for oligomer detection compared to SDS (Fig. 5e). For PFF detection, while Triton X-100 produced higher RAF values, SDS resulted in higher MPR values (Fig. 5f). The relatively lower MPR observed for PFF seeds with Triton X-100 is likely attributable to an excessively high seed concentration, a phenomenon further investigated in Fig. 9. Overall, given its superior aggregation kinetics and reproducibility, and its potential to allow discrimination between oligomer- and PFF-seeded reactions, we selected Triton X-100 as the preferred detergent for α-Syn oligomer detection in the Nano-QuIC assay.

**Figure 5.**
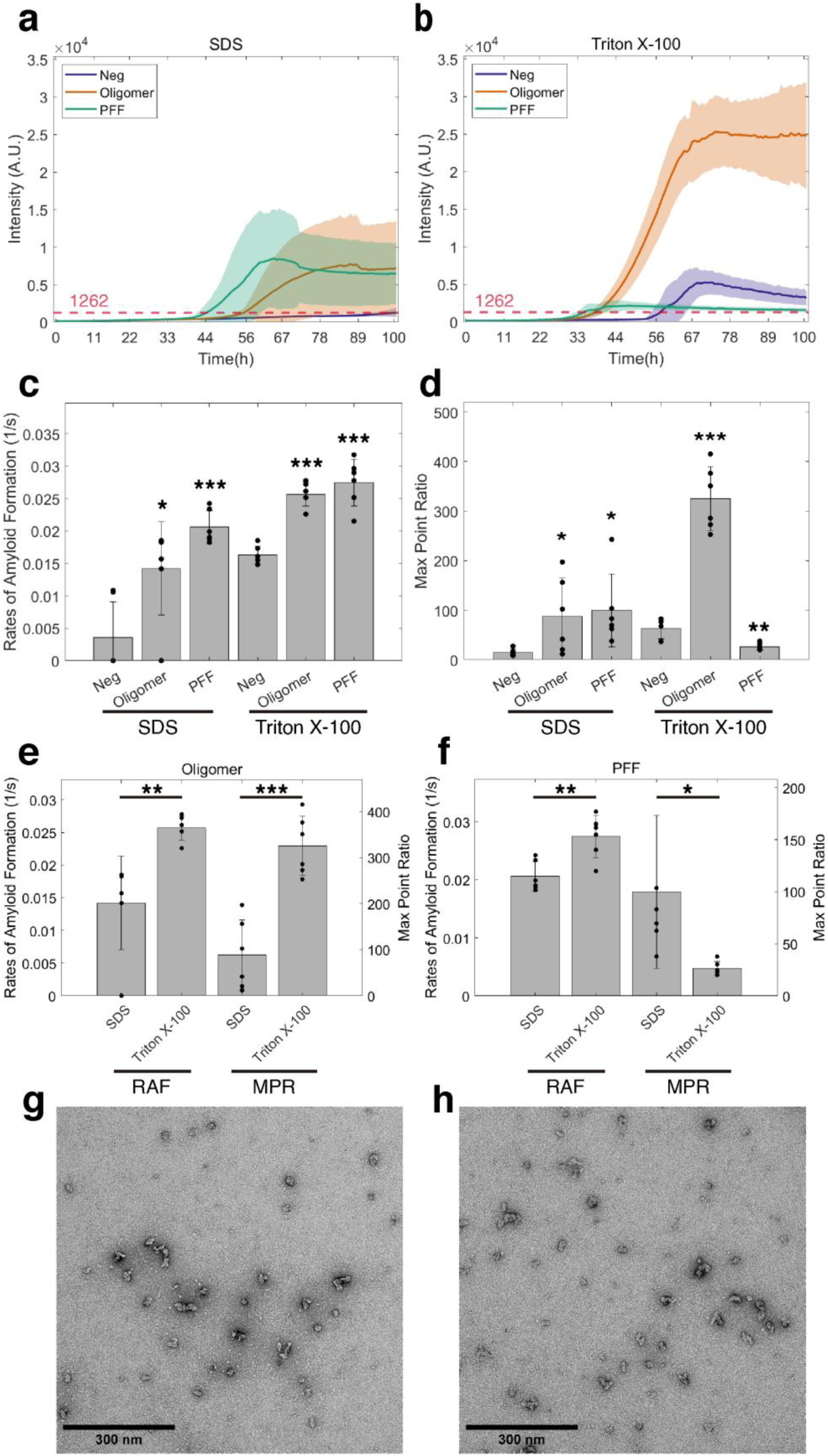
Effect of detergent type on the detection of α-Syn oligomer and PFF seeds. Average ThT fluorescence curves of Nano-QuIC assay with **(a)** 0.002% SDS and **(b)** 0.02% Triton X-100. Solid lines represent the mean, and shaded areas indicate ± SD from six replicates of plasma samples seeded with 100 ng/mL of α-syn oligomer (orange) and 1 μg/mL of PFF seeds (green). The red dashed line indicates the threshold for positive detection. **(c)** RAF and **(d)** MPR of negative controls and seeded plasma samples. Statistical comparisons were made against negative controls. **(e)** RAF and MPR of oligomer seeds and **(f)** RAF and MPR of PFF seeds for each condition. Statistical comparisons were made between SDS and Triton X-100 groups. In all bar graphs, data points represent individual replicates (n = 6), and bars indicate mean ± SD. Statistical significance is denoted as *p < 0.05, **p < 0.01, and ***p < 0.001. TEM images of oligomers at **(g)** 0 hour and **(h)** 48 hours after incubation in the optimized master mix. Scale bars, 300 nm.

To ensure the validity of our results, we assessed the stability of α-Syn oligomer seeds within the optimized master mix. Given the robust amplification observed, it was crucial to confirm whether this activity originated from the oligomers themselves or from spontaneous fibrilization during the lag phase. To investigate this, oligomer seeds were incubated in the optimized master mix (pH 7.4, 500 mM Na^+^, 0.02% Triton X-100) for 48 hours, mimicking the assay duration. TEM imaging at 0 and 48 hours confirmed that the seeds remained in their oligomeric state without undergoing fibrillization (Fig. 5g and 5h). Collectively, these stability results, combined with the superior kinetic performance, confirm the suitability of Triton X-100 for the Nano-QuIC platform.

### Optimized shake cycle for misfolded α-Syn oligomer detection

Next, we improved the shaking protocol for the Nano-QuIC assay. Several studies on various amyloidogenic proteins have shown that longer shaking intervals can accelerate RT-QuIC kinetics (43). To evaluate this effect, we compared two shaking protocols: the standard 60 s shake / 60 s rest (60/60) cycle and a more intensive 100 s shake / 20 s rest (100/20) cycle. The latter increases the shaking duration while shortening the rest time, maintaining a total cycle length of 120 seconds. Both protocols were evaluated in quadruplicate under the previously optimized conditions: pH 7.4, 500 mM Na⁺, 0.02% Triton X-100, and 42°C.

ThT fluorescence curves in Fig. 6a-b show the kinetics of α-Syn aggregation seeded with 100 ng/mL of α-Syn oligomers or 1 μg/mL of PFFs spiked into human plasma. Both the 60/60 and 100/20 shake/rest cycles supported efficient fibril amplification. However, oligomer-seeded reactions exhibited significantly enhanced α-Syn aggregation with the 100/20 protocol. α-Syn aggregation seeded by oligomers typically began at approximately 34 hours with the 60/60 cycle and at approximately 24 hours with the 100/20 cycle. In the case of PFFs, fibril amplification started at approximately 31 hours with the 60/60 cycle and 25 hours with the 100/20 cycle. Both cycles produced some false positives, but the 100/20 cycle exhibited slower and less spontaneous misfolding than the 60/60 cycle (Fig. 6c-d). For oligomer detection, the 100/20 cycle demonstrated significantly higher RAF and MPR values than the 60/60 cycle (Fig. 6e). Similarly, for PFF detection, the 100/20 cycle outperformed the 60/60 cycle in both RAF and MPR (Fig. 6f). Overall, the 100/20 cycle produced stronger and more consistent signals than the 60/60 cycle, suggesting enhanced aggregation kinetics and improved assay consistency across the replicates. Based on these results, we selected the 100/20 cycle as the optimal shaking condition for oligomer detection in the Nano-QuIC assay.

**Figure 6.**
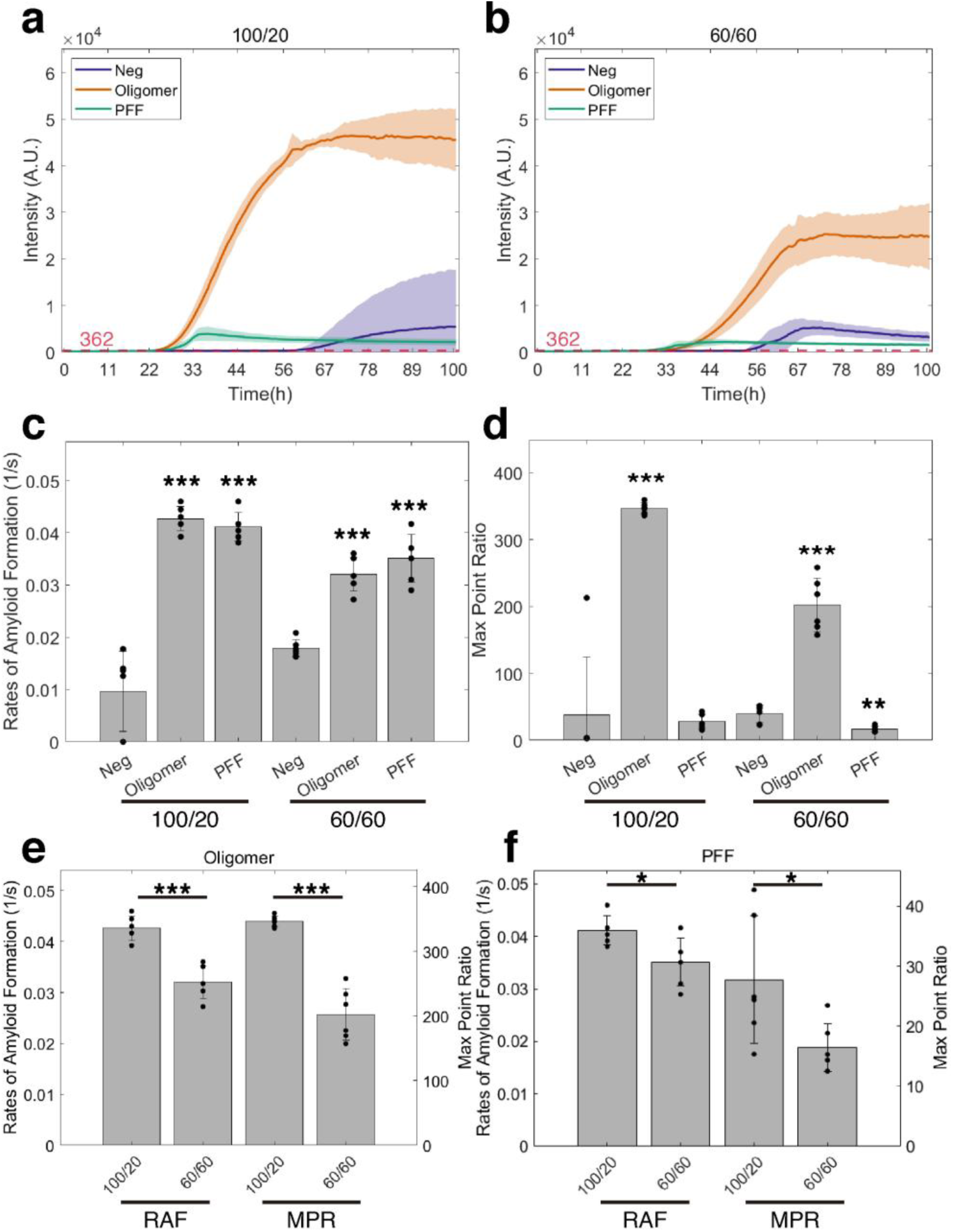
Effect of shake cycles on the detection of α-Syn oligomers and PFF seeds. Average ThT fluorescence curves of Nano-QuIC assay under **(a)** 100/20 and **(b)** 60/60 shake cycles (shake/rest). Solid lines represent the mean, and shaded areas indicate the SD (n = 6) (Data for 60/60 shake cycles are replotted from Fig. 5b for comparison). Samples were seeded with 100 ng/mL of α-syn oligomers (orange) and 1 μg/mL of PFF seeds (green) in human plasma. The red dashed line indicates the threshold for positive detection. **(c)** RAF and **(d)** MPR of negative controls and seeded plasma samples. Statistical comparisons were made against negative controls. **(e)** RAF and MPR of oligomer seeds and **(f)** RAF and MPR of PFF seeds for each condition. Statistical comparisons were made between 100/20 and 60/60 cycles. In all bar graphs, data points represent individual replicates (n = 6), and bars indicate mean ± SD. Statistical significance is denoted as *p < 0.05, **p <0.01, and ***p < 0.001.

### Sensitivity and detection limits of α-Syn oligomers and PFFs under optimized Nano-QuIC conditions

Using the optimized Nano-QuIC assay conditions, we next evaluated the detection limits for α-Syn oligomers and PFFs by performing 10-fold serial dilutions of each seed type spiked into plasma. Each dilution was tested in six replicates to evaluate the assay’s sensitivity. Starting from 100 ng/mL of α-Syn oligomers and 1 μg/mL of PFFs, we performed serial dilutions down to 10^-6^ for oligomers (0.1 pg/mL) and 10^-7^ for PFFs (0.1 pg/mL) under the optimized condition: pH 7.4, 500 mM Na⁺, 0.02% Triton X-100, and 100/20 s shake/rest cycle at 42°C.

As shown in Fig. 7a-d, S2 and S3, α-Syn oligomer seeds were reliably detected down to 100 pg/mL, and PFF seeds were detectable at concentrations as low as 10 pg/mL. Detection was defined as ThT fluorescence exceeding the established threshold in at least 50% of replicates per condition within 150 hours. These results highlight the assay’s capability to detect α-Syn seeds at low pg/mL concentrations, demonstrating its potential for high-sensitivity diagnostic applications in clinical matrices.

**Figure 7.**
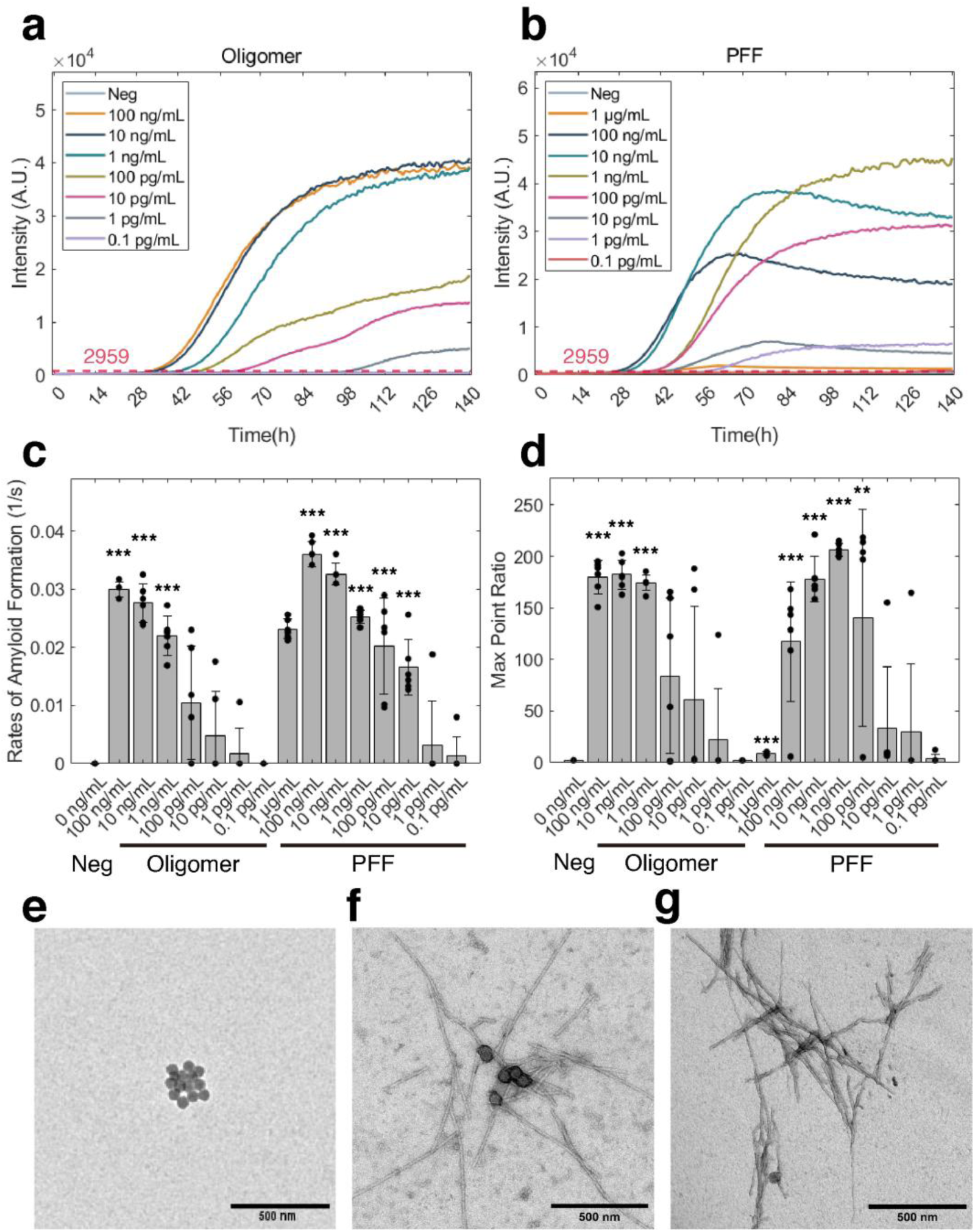
Detection sensitivity of Nano-QuIC for α-Syn oligomers and PFFs across serial dilutions. Average ThT fluorescence curves of Nano-QuIC assays performed on plasma spiked with various concentrations of **(a)** α-Syn oligomers and **(b)** PFFs. Solid lines represent the mean of six replicates. Seed concentrations spiked into human plasma range from 100 ng/mL to 0.1 pg/mL for oligomers and 1 μg/mL to 0.1 pg/mL for PFFs. **(c)** RAF and **(d)** MPR of negative controls and seeded plasma samples. Statistical comparisons were made against negative controls. In bar graphs, data points represent individual replicates (n = 6), and bars indicate mean ± SD. Statistical significance is denoted as *p < 0.05, **p < 0.01, and ***p < 0.001. Representative TEM images of reaction products collected at 88.5 hours for **(e)** unseeded control, **(f)** oligomer-seeded, and **(g)** PFF-seeded samples. Scale bars, 500 nm.

As briefly mentioned in the optimized detergent section, excessive fibril seed concentration appears to inhibit assay performance compared to lower concentrations (Fig. 7b and 7d). This inhibition is possibly driven by the excessive aggregation and entanglement of fibrils and siNPs (Fig. S4). These dense complexes may persist despite mechanical agitation, thereby diminishing the availability of accessible growth sites and reducing the number of active siNPs. To further validate the reaction products, we performed TEM imaging on Nano-QuIC samples with no seed, oligomer seeds, or PFF seeds. As shown in Fig. 7e, reaction products obtained in the absence of seeds showed no fibril formation, with only siNPs visible. In contrast, the addition of either α-Syn oligomers or PFFs resulted in the clear formation of fibrils associated with the siNPs (Fig. 7f and 8g). These results confirm that both seed species successfully initiate α-Syn aggregation within the Nano-QuIC platform, validating the kinetic signals observed.

## Discussion

In this study, we optimized the Nano-QuIC assay to achieve ultrasensitive detection of α-Syn oligomers in human plasma. This represents a critical step toward developing a minimally invasive diagnostic tool for early-stage PD. The focus on oligomers is crucial; while their exact pathogenic mechanisms remain under investigation, there is a broad consensus that oligomer species induce cellular damage by disrupting membranes, impairing mitochondrial function and protein degradation pathways, damaging synapses, and triggering inflammatory responses, leading to neurodegenerative diseases. (6–10,23,44,45). They can also propagate pathology between cells in a prion-like manner (10).

Furthermore, the significance of oligomers is underscored by recent studies of *LRRK2* mutation carriers who exhibit typical Parkinsonian symptoms despite the absence of classical Lewy body pathology (13,14). This clinical paradox has been elucidated by proximity ligation assays (PLA), which revealed that these Lewy body-negative cases harbor high levels of α-Syn oligomers. Such findings suggest that oligomer accumulation may contribute significantly to neurodegeneration even in the absence of extensive fibrillization, highlighting the critical role of these species in the pathogenic process of PD.

Despite their clinical relevance, the primary barrier to using oligomers as biomarkers has been their exceedingly low concentrations in accessible biological fluids like blood. The development of SAAs, including RT-QuIC, has been a major breakthrough for detecting misfolded α-Syn (31,32). However, its application has been most successful with post-mortem tissue or invasively obtained CSF. While CSF is valuable due to its proximity to the CNS, the requirement for lumbar punctures limits its utility for routine screening. In contrast, plasma has emerged as an attractive alternative because it can be collected less invasively and contains detectable levels of α-Syn (20–22,34). Our previous study demonstrated that the incorporation of siNPs into the conventional RT-QuIC assay enhances the detection of α-Syn fibrils by mitigating the effects of inhibitory components commonly present in complex human biological fluids (35).

Building on this concept, we first compared the performance of the standard RT-QuIC and Nano-QuIC assays using α-Syn oligomer and PFF seeds spiked into human plasma. Our comparative analysis revealed distinct advantages of the Nano-QuIC format. While Nano-QuIC demonstrated greater robustness and consistency in detecting PFFs compared to standard RT-QuIC, its most striking advantage was the unique capability to detect α-Syn oligomers, which were undetectable by the standard assay. We subsequently optimized Nano-QuIC assay conditions to maximize performance in detecting α-Syn oligomers in plasma samples. Through systematic evaluation of buffer pH, Na⁺ concentrations, detergent types, and shaking cycles, we established an optimal protocol: pH 7.4 using PBS buffer, 500 mM total Na⁺ concentration, 0.02% Triton X-100, 100 s shaking followed by 20 s rest cycles, at an incubation temperature of 42°C.

Under these optimized conditions, we assessed the assay’s sensitivity by performing 10-fold serial dilutions of oligomer and PFF seeds. Oligomers were reliably detected at concentrations as low as 100 pg/mL, while PFFs were detectable down to 10 pg/mL. These results underscore the assay’s capability to detect trace pathological seeds, a prerequisite for clinical application. For comparison, our previous Nano-QuIC study, which detected α-Syn PFFs spiked into human plasma, reported a detection limit of 900 pg/mL (35). This current study further highlights the strong potential of Nano-QuIC assays as a highly sensitive platform for detecting α-Syn aggregates in blood-based diagnostics.

The exceptional sensitivity of the Nano-QuIC assay in complex biofluids like plasma is primarily attributed to the electrostatic and hydrophobic interactions between siNPs and the target proteins (35). This surface-mediated mechanism enables α-Syn species to effectively sequester from inhibitors and concentrate on the siNP surfaces, leading to a marked increase in the local concentration of both seeds and substrates. This mechanistic model is further corroborated by TEM imaging of the reaction products (Fig. 7f and 7g), which clearly shows newly formed fibrils sprouting and elongating directly from the siNPs.

Notably, we observed a reduction in assay efficiency at excessively high PFF concentrations, a phenomenon possibly driven by the rapid and dense growth of fibrils on siNP surfaces. This accelerated growth may promote the formation of large, entangled complexes where fibrils bridge multiple siNP assemblies (Fig. S4). Such entanglement may physically restrict access to active seeding sites and reduce the mobility of siNPs, thereby limiting further amplification. In contrast, this effect was not observed in oligomer-seeded reactions within the tested range. This is likely due to the inherently slower aggregation kinetics of oligomers, resulting in smaller, more sparsely distributed fibril-siNP complexes that remain susceptible to fragmentation by mechanical agitation. These morphological characteristics provide a structural basis for the consistent performance of the Nano-QuIC assay across a broad range of oligomer concentrations.

While this study was conducted using synthetic α-syn seeds, which may not fully recapitulate the structural heterogeneity and complexity of endogenous aggregates, the sensitivity achieved here is promising for clinical translation. As endogenous oligomer levels in patients are expected to be extremely low (46–48), the capacity to detect seeds in the low picogram range is essential. Furthermore, integrating our assay with recent methods such as the Constant Shake-Induced Conversion (CSIC) assay (34) could potentially enhance the detection of oligomers in early-stage PD. Future work should focus on validating this assay in large, well-characterized clinical cohorts to assess diagnostic accuracy and correlate oligomer burden with disease progression (49).

Notably, this platform could be instrumental in identifying patients whose pathology is primarily driven by oligomeric species, such as certain *LRRK2* variants. By ensuring the detection of the oligomeric fraction that standard assays often miss, our assay provides a powerful tool to more accurately represent the total pathological burden in blood. This capacity is essential for stratifying patients for clinical trials and ensuring that therapeutic interventions are tailored to the specific proteinopathy of the individual. By enabling the detection of oligomeric species, which likely precede fibril formation, this optimized Nano-QuIC platform offers a viable path toward a non-invasive, early diagnostic tool for PD.

## Materials and Methods

### QuIC amplification of α-Syn

All RT-QuIC and Nano-QuIC reactions were performed using black, clear-bottom, 96-well microplates (Thermo Scientific). To optimize the assay for α-syn oligomers and PFFs, several reaction conditions were tested. For the initial RT-QuIC analysis, the master mix contained 1× phosphate buffer (9.2 mM NaH₂PO₄, 2.8 mM Na₂HPO₄, 2.7 mM KCl, 137 mM NaCl), 1 mM ethylenediaminetetraacetic acid (EDTA), 170 mM NaCl (in addition to the NaCl present in the phosphate buffer), 10 μM Thioflavin T (ThT), and 0.09 mg/mL recombinant α-Syn monomers (ND BioScience, Epalinges, Switzerland). For the Nano-QuIC assay, 2.5 mg/mL of 50 nm silica nanoparticles (Fortis Life Sciences Company) were added to the same master mix.

A volume of 98 μL of master mix was dispensed into each well, followed by the addition of 2 μL sample, resulting in a total volume of 100 μL per well. After sealing, the plates were incubated in a VANTAstar plate reader (BMG Labtech, Germany) at 42°C for at least 150 hours with shaking at 700 rpm (double orbital mode; 60 s shaking followed by 60 s rest). All solutions were filtered through 0.22 μm sterile filters prior to use. Recombinant α-Syn monomers were purified by centrifugation at 3000 × g for 15 minutes at room temperature using a 100 kDa MWCO centrifugal filter (Pall Corporation) before being added to the reaction. To further optimize assay conditions for α-Syn oligomer detection, we tested various parameters, including pH, Na⁺ concentrations, detergent types, and shaking protocols. The optimized master mix was modified to contain 1× PBS, 1 mM EDTA, 288 mM NaCl, 10 μM ThT, 2.5 mg/mL siNPs, and 0.09 mg/mL recombinant α-Syn monomers. Plates were incubated at 42°C for at least 150 hours with shaking at 700 rpm (double orbital mode; 100 s shaking followed by 20 s rest). Fluorescence intensity, expressed in relative fluorescence units (RFU), was measured every 45 minutes using an excitation wavelength of 450 ± 10 nm and an emission wavelength of 480 ± 10 nm, with bottom reading and a gain setting of 515.

### Preparation of Parkinson’s disease (PD) mimicking plasma samples

Alpha-synuclein (a-Syn) monomers, oligomers, and preformed fibrils (PFFs) were purchased from ND BioSciences (Epalinges, Switzerland). All a-Syn species were of the wild-type (WT) form and were characterized by transmission electron microscopy (TEM), SDS-PAGE, protein concentration estimation, mass spectrometry, and far-UV circular dichroism (CD) spectroscopy to confirm their structural properties by ND BioSciences (Supplementary Datasheet S1). Prior to use, proteins were aliquoted and stored at −80°C and working concentrations were prepared in phosphate-buffered saline (1× PBS, pH 7.4) immediately before each experiment. Healthy human EDTA plasma sample was purchased from Innovative research, Inc (Novi, MI, USA). All samples were stored at −80°C until use and thawed on ice prior to analysis. To prepare PD-mimicking plasma samples, α-Syn oligomers or PFFs were first diluted in 1× PBS to the desired concentrations. A volume of 40 μL of the diluted oligomer or PFF solution was then spiked into 360 μL of healthy control (HC) human plasma, resulting in a total volume of 400 μL. The spiked plasma samples were transferred into 1.5 mL microcentrifuge tubes and centrifuged at 21,000 × g for 40 minutes at 4°C. Following centrifugation, the supernatant was carefully aspirated, and the resulting pellet was resuspended in 1% Triton X-100. A 2 μL aliquot of the resuspended pellet was then added to each well containing 98 μL of the RT-QuIC or Nano-QuIC master mix, yielding a final volume of 100 μL per reaction.

### Transmission electron microscopy

TEM samples were prepared using Carbon-coated, 400-mesh copper grids (Electron Microscopy Sciences). The grids were first discharged with plasma by easiGlow discharge system (PELCO) for 60 seconds to enhance sample adhesion. The samples were then fixed with a fixative buffer (2% glutaraldehyde, 0.1 M Cacodylate in water), followed by negative staining with 2% (w/v) uranyl acetate. Each incubation step was performed at room temperature and included a washing step with deionized (DI) water. Excess solution and DI water were carefully removed from the grid using filter paper after each step. The grids were examined using a Tecnai Spirit Bio-Twin Transmission Electron Microscope (Field Electron and Ion Company) equipped with a 2k × 2k CCD camera. TEM imaging was performed at the University of Minnesota’s Characterization Facility. The sizes of the oligomers and the lengths of the PFFs were measured using ImageJ (50). The size of each oligomer was determined using the equivalent circular radius.

### Kinetic data analysis

The rate of amyloid formation (RAF) was calculated as the inverse of the time (s⁻¹) required for the fluorescence signal to reach the threshold value. The fluorescence threshold was set as 30 times the standard deviation of the baseline signal, with the baseline defined as the mean fluorescence intensity at cycle 3 across all replicates. The maximum point ratio (MPR) was calculated by dividing the peak fluorescence intensity of each replicate by its baseline fluorescence value (36). Statistical differences in RAF and MPR between case and negative control groups were evaluated using a t-test. Data variability is presented as mean ± SD, and statistical significance was defined as p < 0.05. All kinetic analyses were performed using custom scripts written in MATLAB (MathWorks), with reference to QuICSeedR, an R package developed for RT-QuIC assay analysis (51).

## Supporting information

Supplementary datasheet S1

## Author contributions

H.L, P.A.L., S.-H.O., and H.Y.P. conceived and supervised the research. H.J., P.R.C., and H.A. performed experiments. H.L. provided alpha synuclein oligomer and fibrils. All authors reviewed and analyzed the results. All authors wrote and contributed to the final manuscript.

## Acknowledgements

We thank Manci Li for sharing her R code and providing guidance on its use. We thank Fang Zhou for assistance with TEM imaging. H.J., P.R.C., H.A., P.A.L., S.-H.O., and H.Y.P. were supported by the Minnesota Partnership for Biotechnology and Medical Genomics grant from the State of Minnesota for the establishment of the Center for Advanced Synucleinopathy Diagnostics (ASCEND). P.R.C. and S.-H.O. acknowledge support from the Michael J. Fox Foundation for Parkinson’s Research (Seed Amplification Assay Innovation Program). H.Y.P. acknowledges the startup funding from the University of Minnesota. S.-H.O. further acknowledges support from the Sanford P. Bordeau Chair and the International Institute for Biosensing (IIB) at the University of Minnesota.

## Competing interests

P.R.C., P.A.L., and S.-H.O. are co-inventors on a patent related to the Nano-QuIC assay. P.A.L. and S.-H.O. are co-founders and equity holders of Priogen Corp., a diagnostic company specializing in the detection for prions and protein misfolding diseases. H.A.L. is a co-founder and chief scientific officer of ND BioSciences SA, a company focusing on developing novel therapies and diagnostics for neurodegenerative diseases. All other authors declare no competing interests.

