## Supplementary Information for "Ultrasensitive Detection of Alpha-Synuclein Oligomers in Human Plasma Using Optimized Nano-QuIC"

[3]ND BioSciences SA, Switzerland.

[4]Weill Cornell Medicine Qatar, Education City, Qatar Foundation, Doha, Qatar.

[5]Minnesota Center for Prion Research and Outreach, University of Minnesota, St. Paul, Minnesota 55108, United States.

[6]Department of Veterinary and Biomedical Sciences, University of Minnesota, St. Paul, Minnesota 55108, United States.

*Corresponding authors:

Sang-Hyun Oh, Hye Yoon Park

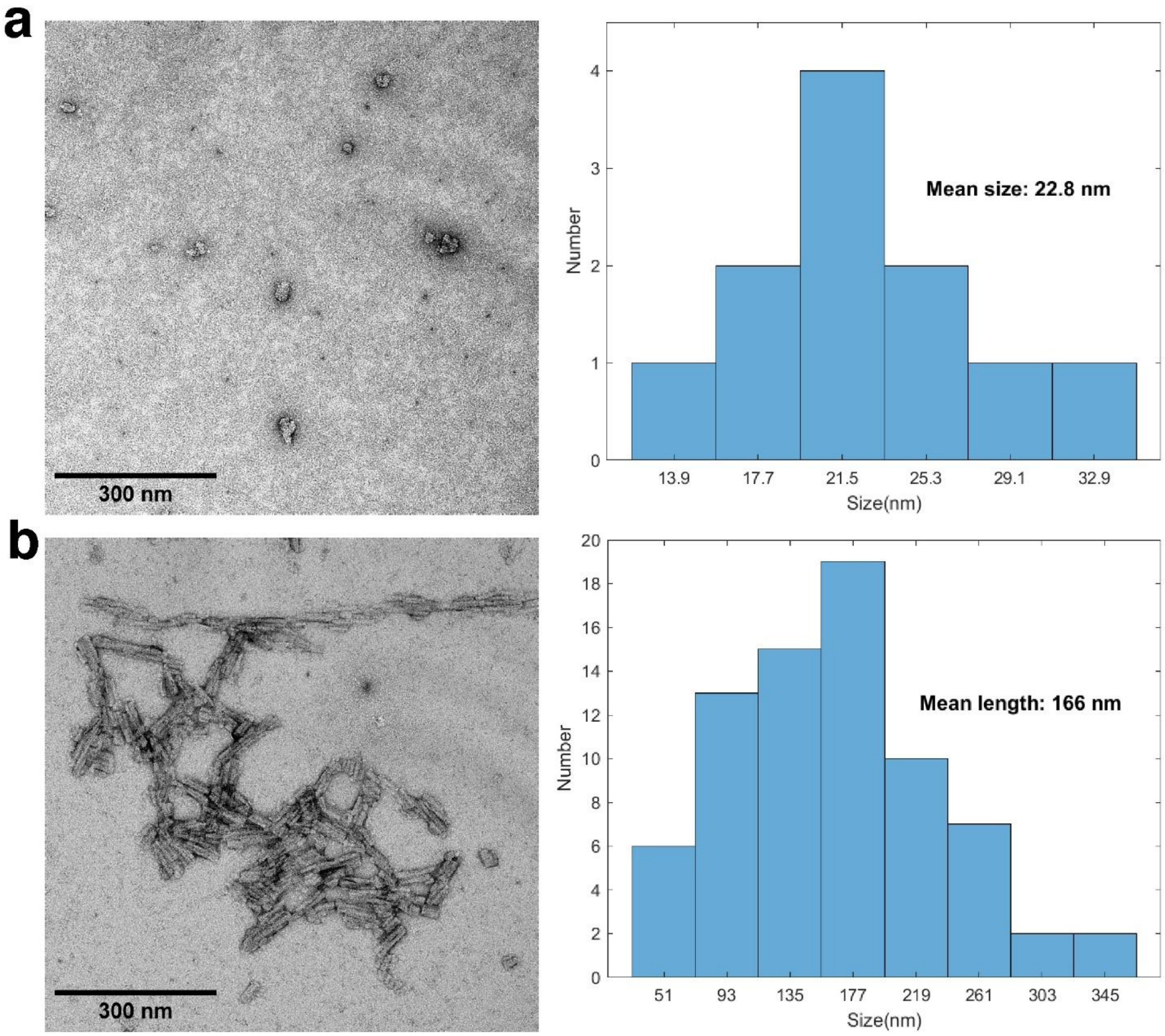


**Figure S1. Morphological characterization of α-Syn seeds.** Representative TEM images and corresponding size distributions of **(a)** α-Syn oligomers and **(b)** PFFs in PBS buffer. Scale bars, 300 nm.

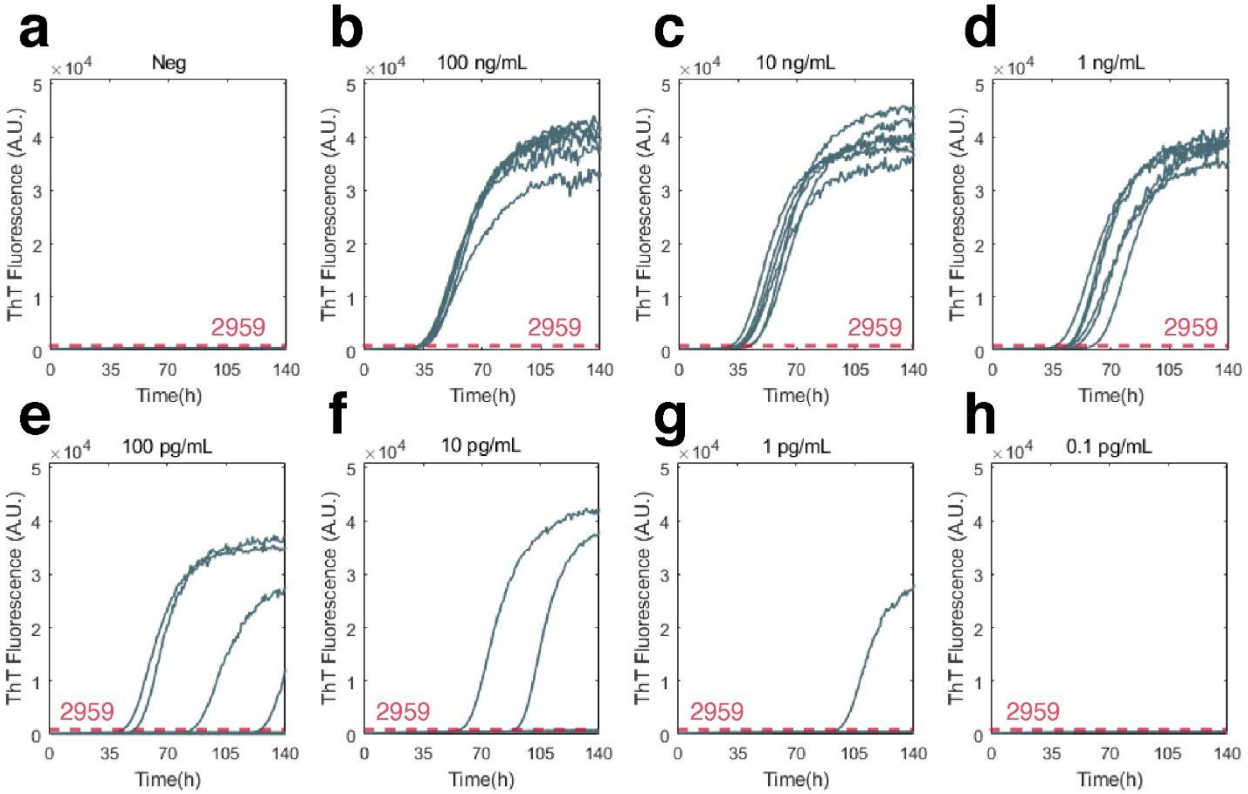


**Figure S2. Individual ThT fluorescence curves of Nano-QuIC assays seeded with varying concentrations of α-Syn oligomers.** Solid lines represent individual traces from six replicate reactions for each condition: **(a)** Negative control (unseeded), **(b)** 100 ng/mL, **(c)** 10 ng/mL, **(d)** 1 ng/mL, **(e)** 100 pg/mL**, (f)** 10 pg/mL, **(g)** 1 pg/mL, and **(h)** 0.1 pg/mL.

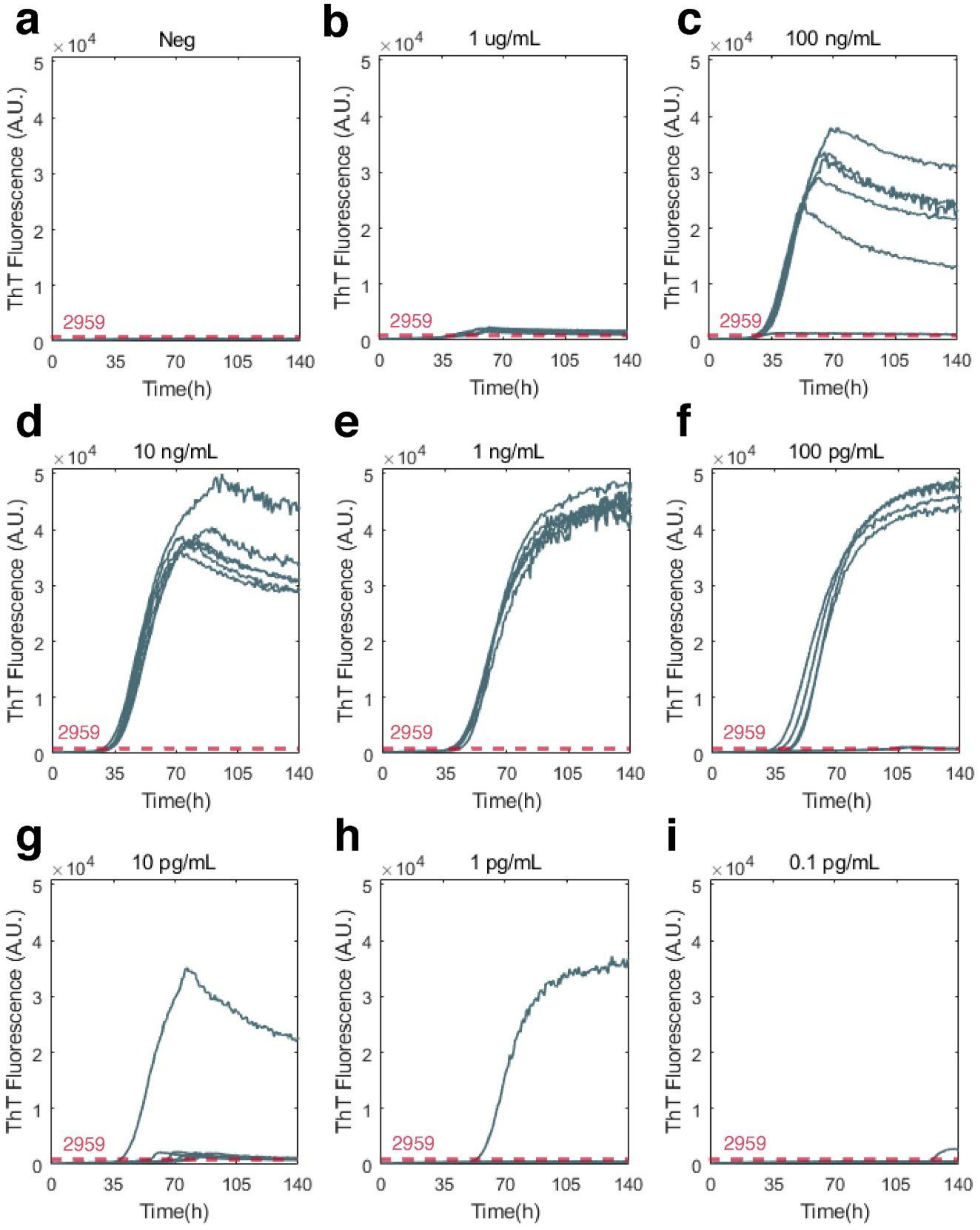


**Figure S3. Individual ThT fluorescence curves of Nano-QuIC assays seeded with varying concentrations of α-Syn PFF.** Solid lines represent individual traces from six replicate reactions for each condition: **(a)** Negative control (unseeded), **(b)** 1 μg/mL, **(c)** 100 ng/mL, **(d)** 10 ng/mL, **(e)** 1 ng/mL**, (f)** 100 pg/mL, **(g)** 10 pg/mL, **(h)** 1 pg/mL, and **(i)** 0.1 pg/mL.

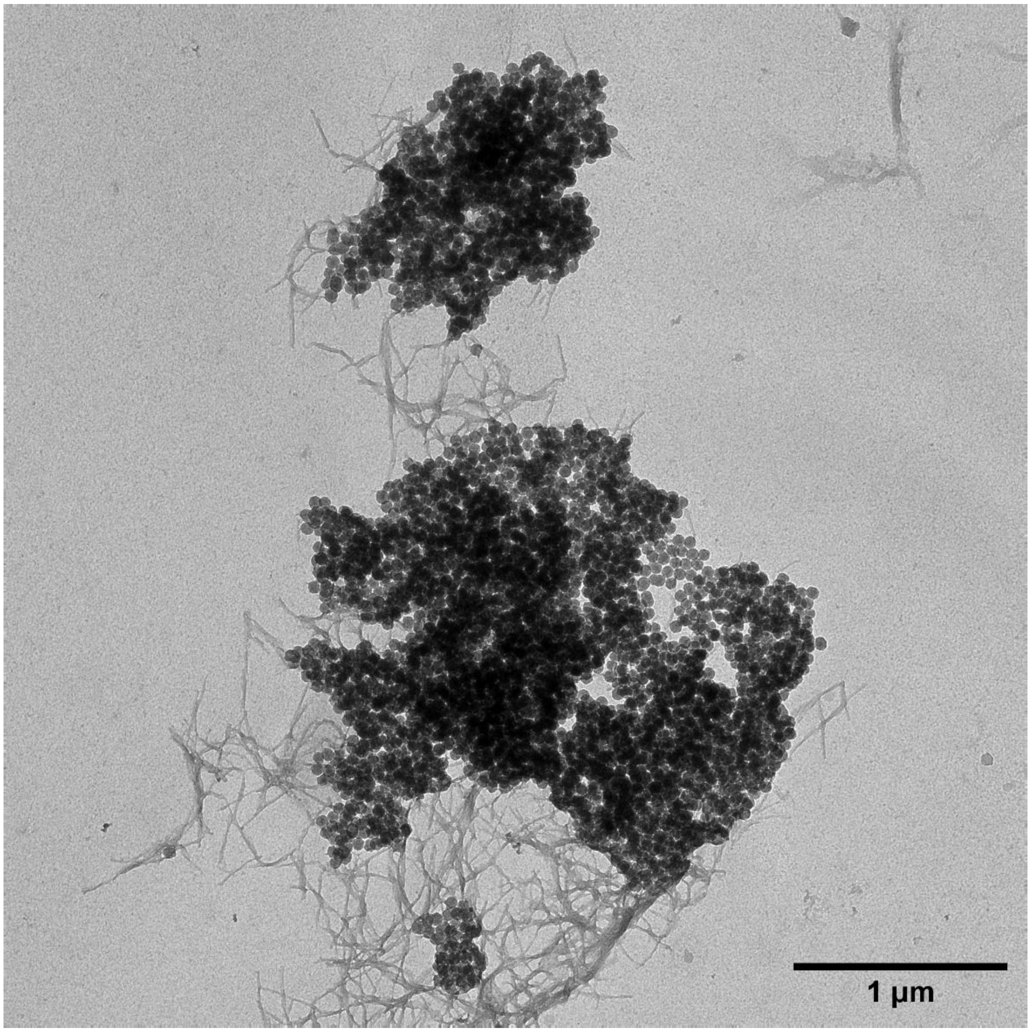


**Figure S4. Dense fibril growth and large entangled complex formation on siNP surfaces.** A representative TEM image displaying fibril-mediated bridging and the entanglement of multiple siNPs. Scale bar, 1 μm.
