## Supplementary datasheet S1 for "Ultrasensitive Detection of Alpha-Synuclein Oligomers in Human Plasma Using Optimized Nano-QuIC"

BIOSCIENCES

### Product Datasheet

#### Human α-Synuclein Monomers

| | |
|---|---|
| **Sequence** | MDVFMKGLSKAKEGVVAAAEKTKQGVAEAAGKTKEGVL YVGSKTKEGVVHGVATVAEKTKEQVTNVGGAVVTGVTA VAQKTVEGAGSIAAATGFVKKDQLGKNEEGAPQEGILED MPVDPDNEAYEMPSEEGYQDYEPEA |
| **Swiss Prot** | P37840 |
| **Gene ID** | 6622 |
| **Accession #** | NP_000336.1 |
| **Species** | Human |
| **Amino acids** | 1-140, full length protein |
| **Conjugates/Tags** | No Tag |
| **Molecular weight** | 14 kDa (14,460 Da) |
| **Nature** | Recombinant, expressed in Escherichia coli |
| **Certificate of analysis** | Certified > 95 % SDS-PAGE. Full characterization provided in Figure 1. |
| **Field of Use** | Not for use in humans. For research purposes only. |
| **Applications** | In vitro assays, cellular assays, animal studies or as standards in WB, SDS-PAGE, ELISA, and other immunoassays. |
| **Form** | Shipped lyophilized on dry ice. |
| **Preparation** | Resuspend in appropriate buffer for downstream assays. We recommend resuspension in PBS, at a concentration range 0.1 – 2 µg/µl. |
| **Storage** | Store at -80°C upon receipt. Following resuspension, aliquot and store at -80°C. |
| **Handling** | We recommend avoiding repeated freeze-thaw cycles. |
| **Product Citation** | Please cite this product as "Human α-Synuclein Monomers (ND Biosciences SA, Switzerland, Catalogue #ND001 – Lot #07/22-001.001.2)" |
| **Safety measures** | This product is an active protein and may elicit a biological response in vivo, handle with caution. |
| **References** | Detailed protocols on how to handle this protein and remove any preformed aggregates are presented in: Kumar ST, Donzelli S, Chiki A, Syed MMK, Lashuel HA. A simple, versatile and robust centrifugation-based filtration protocol for the isolation and quantification of α-synuclein monomers, oligomers and fibrils: Towards improving experimental reproducibility in α-synuclein research. J Neurochem. 2020;153(1):103-119. doi:10.1111/jnc.14955. |

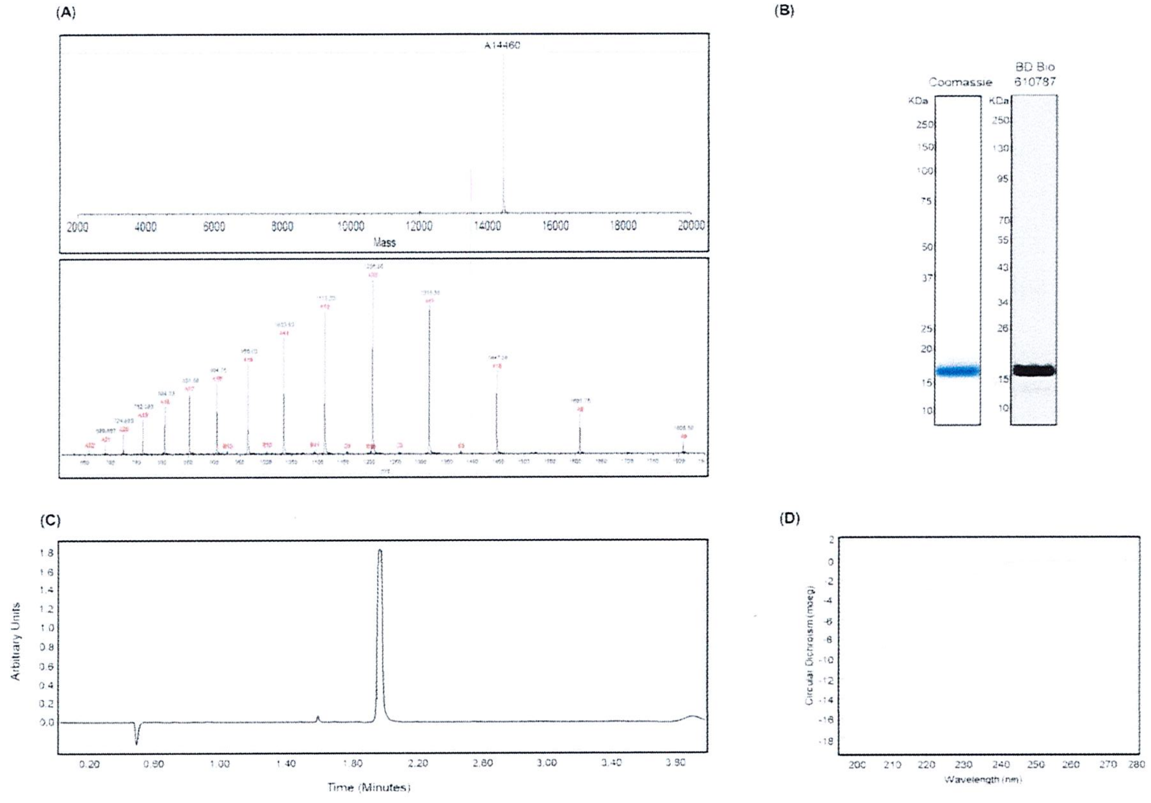


**Figure 1. Characterization of α-Syn monomers. (A)** Mass Spec (ESI) analysis confirms the integrity of the α-Syn protein and shows a mass of ~14460 Da. **(B)** Coomassie staining (left) at 1 µg loaded amount shows that α-Syn migrates as a single band at ~14 kDa. Western blotting (right) confirms reactivity of α-Syn monomers (at 100ng loaded amounts) with the following antibody: BD-Biosciences 610787, epitope: 91-99. **(C)** UPLC analysis shows a single peak at ~2min, establishing the purity of the protein. **(D)** CD analysis shows a spectrum with a minimum at ~200 nm, establishing disordered conformational ensemble of α-Syn monomers.

BIOSCIENCES

### Product Datasheet

#### Unmodified Human α-Synuclein Oligomers

| | |
|---|---|
| **Sequence** | MDVFMKGLSKAKEGVVAAAEKTKQGVAEAAGKTKEGVL YVGSKTKEGVVHGVATVAEKTKEQVTNVGGAVVTGVTA VAQKTVEGAGSIAAATGFVKKDQLGKNEEGAPQEGILED MPVDPDNEAYEMPSEEGYQDYEPEA |
| **Swiss Prot** | P37840 |
| **Gene ID** | 6622 |
| **Accession #** | NP_000336.1 |
| **Species** | Human |
| **Amino acids** | 1-140, full length protein |
| **Conjugates/Tags** | No Tag |
| **Molecular weight** | 14 kDa (14,460 Da) |
| **Nature** | Recombinant, expressed in Escherichia coli |
| **Certificate of analysis** | Certified > 95 % purity by SDS-PAGE. Full characterization provided in Figure 1. |
| **Field of Use** | Not for use in humans. For research purposes only. |
| **Applications** | In vitro assays, cellular assays, animal studies or as standards in WB, SDS-PAGE, ELISA, and other immunoassays. |
| **Form** | Shipped on dry ice. |
| **Preparation** | Protein is prepared in PBS at 0.12 mg/ml. |
| **Storage** | Store at -80°C upon receipt. |
| **Handling** | Protein is stable for up to 3 freeze/thaw cycles. We recommend avoiding repeated thawing cycles. |
| **Product Citation** | In case of publication or scientific presentations using this product, please cite as “Unmodified Human α-Synuclein Oligomers (ND Biosciences SA, Switzerland, Catalogue #ND002, Lot #11/21-002.001.1)”. Characterization data (Figure 1) remains property of ND Biosciences and is not to be used in any publications without written permission from ND Biosciences. |
| **Safety measures** | This product is an active protein and may elicit a biological response in vivo, handle with caution. |
| **References** | Detailed protocols on how to handle this protein are presented in: Kumar ST, Donzelli S, Chiki A, Syed MMK, Lashuel HA. A simple, versatile and robust centrifugation-based filtration protocol for the isolation and quantification of α-synuclein monomers, oligomers and fibrils: Towards improving experimental reproducibility in α-synuclein research. J Neurochem. 2020;153(1):103-119. doi:10.1111/jnc.14955. |

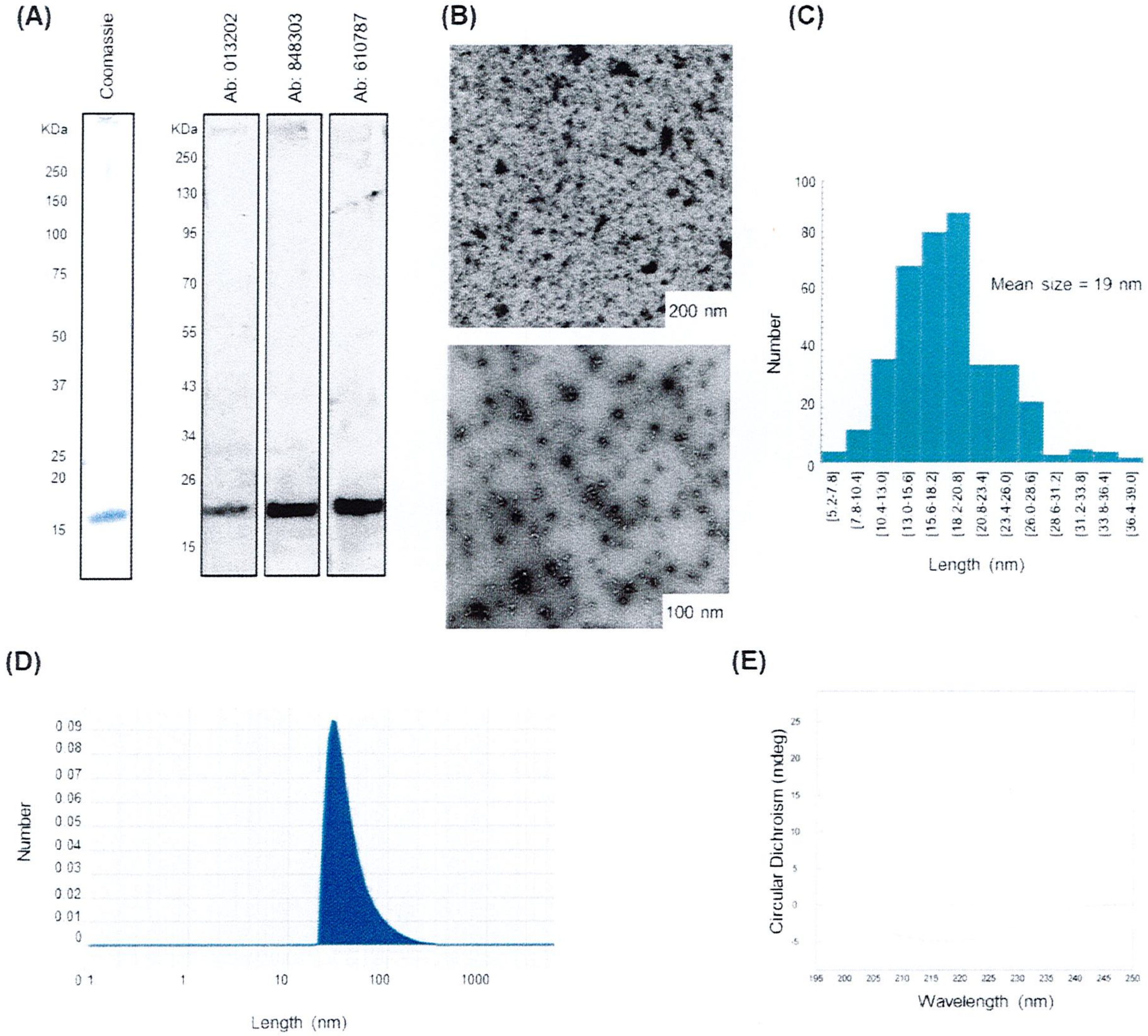


**Figure 1. Characterization of α-Syn oligomers. (A)** Coomassie staining and western blot analysis using different antibodies (Ab: 013202, Abbexa, epitope: 15-64, Ab: 848303, Biolegend, epitope: 80-96, and Ab: 610787, BD-Biosciences, epitope: 91-99) shows that α-Syn oligomers migrate as SDS-resistant high molecular weight (HMW) species (>250kDa), with some oligomers breaking down to monomers at ~14 kDa. **(B)** Transmission electron microscopy (TEM) of uranyl acetate stained α-Syn oligomers shows a population of annular pores and spherical/tubular-like particles. **(C)** The size distribution of oligomers visualized by TEM is between 5 and 39 nm, with an average size of 19 nm. **(D)** Dynamic light scattering analysis of α-Syn monomers shows a homogenous population in terms of size distribution, with average size of ~30 nm. **(E)** CD analysis shows a spectrum with a minimum at 215 nm, establishing β-sheet conformation of α-Syn oligomers.

### Product Datasheet

#### Human α-Synuclein Fibrils

| | |
|---|---|
| **Sequence** | MDVFMKGLSKAKEGVVAAAEKTKQGVAEAAGKTKEGVL YVGSKTKEGVVHGVATVAEKTKEQVTNVGGAVVTGVTA VAQKTVEGAGSIAAATGFVKKDQLGKNEEGAPQEGILED MPVDPDNEAYEMPSEEGYQDYEPEA |
| **Swiss Prot** | P37840 |
| **Gene ID** | 6622 |
| **Accession #** | NP_000336.1 |
| **Species** | Human |
| **Amino acids** | 1-140, full length protein |
| **Conjugates/Tags** | No Tag |
| **Molecular weight** | 14 kDa (14,460 Da) |
| **Nature** | Recombinant, expressed in Escherichia coli. |
| **Certificate of analysis** | Certified > 95 % purity by SDS-PAGE. Full characterization provided in Figure 1. |
| **Field of Use** | Not for use in humans. For research purposes only. |
| **Applications** | In vitro assays, cellular assays, animal studies or as standards in WB, SDS-PAGE, ELISA, and other immunoassays. |
| **Form** | Shipped in PBS solution on dry ice. |
| **Preparation** | Protein was dissolved in PBS at 2 mg/ml. |
| **Storage** | Store at -80°C upon receipt. Following resuspension, aliquot and store at -80°C. |
| **Handling** | Protein is stable for up to 3 freeze/thaw cycles. We recommend avoiding repeated thawing cycles. |
| **Product Citation** | In case of publication or scientific presentations using this product, please cite as Human α-Synuclein Fibrils Polymorph 1 (ND Biosciences SA, Switzerland, Catalogue #ND003, Lot # 05/21-003.002)". |
| **Safety measures** | This product is an active protein and may elicit a biological response in vivo, handle with caution. |
| **References** | Kumar ST, Donzelli S, Chiki A, Syed MMK, Lashuel HA. A simple, versatile and robust centrifugation-based filtration protocol for the isolation and quantification of α-synuclein monomers, oligomers and fibrils: Towards improving experimental reproducibility in α-synuclein research. J Neurochem. 2020;153(1):103-119. doi:10.1111/jnc.14955. |

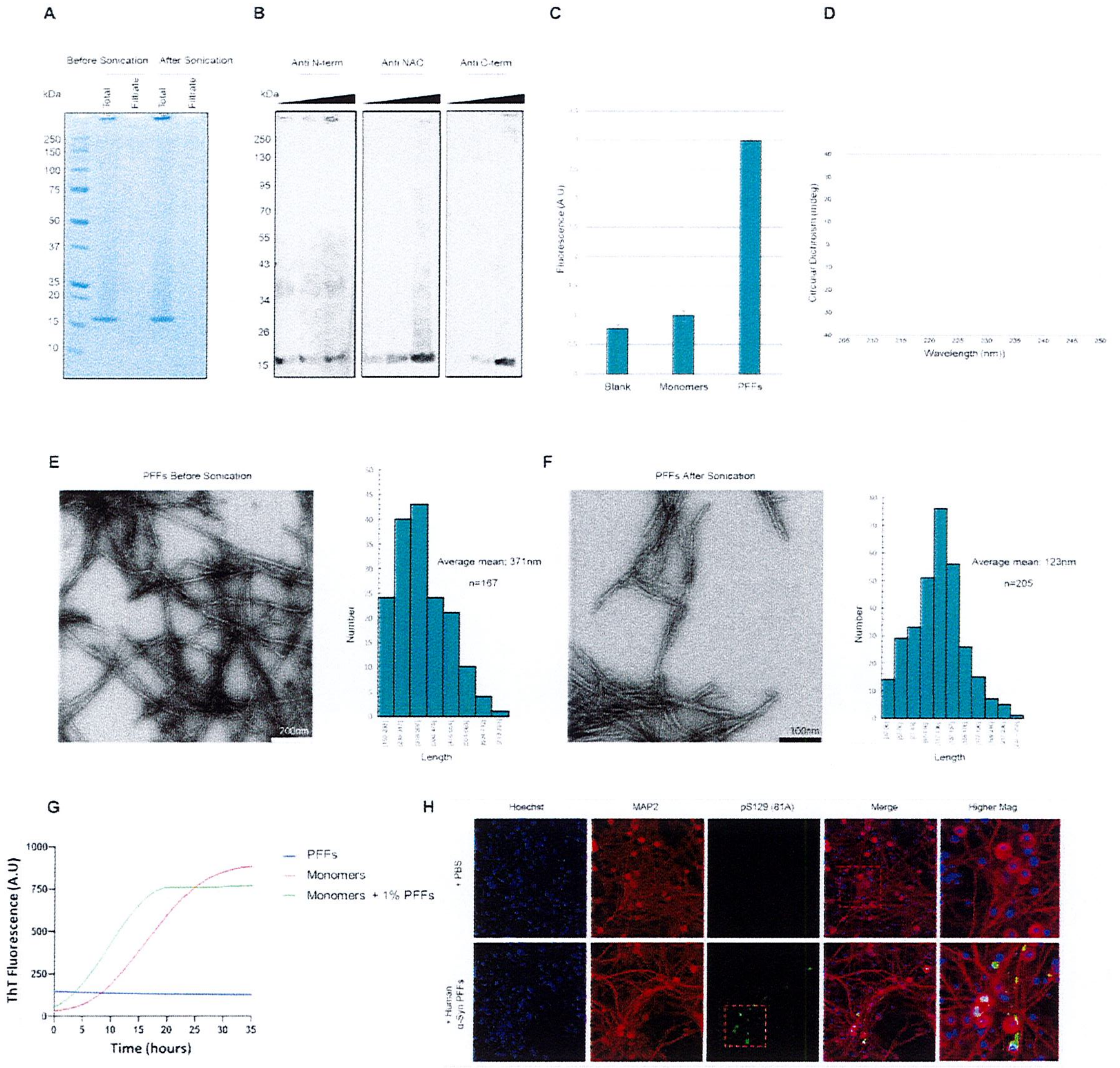


**Figure 1. Characterization of α-Syn preformed fibrils (PFFs). (A)** Coomassie staining shows that α-Syn PFFs encompass very low amount of released soluble monomers (flow through 100 kDa filter) before and after sonication. Coomassie staining also shows that the α-Syn PFFs migrate as SDS-resistant high molecular weight (HMW) species (in the stacking gel, >250kDa), with some fibrils breaking down into monomers at ~14 kDa. **(B)** Screening of reactivity of α-Syn fibrils with a panel of antibodies targeting the N-terminus, NAC region and C terminus of α-Syn at 3 different loaded concentrations (50ng,100ng,200ng). Antibodies used: Abbexa (Abx008695), BioLegend (848302, epitope: 80-96), BD-Biosciences (610787, epitope 91-99) **(C)** Assessment of ThioflavinT (ThT) binding shows that α-Syn PFFs after sonication binds to ThT and comprise amyloid structure, unlike the blank and α-Syn monomer controls. **(D)** Circular dichroism (CD) analysis shows a spectrum with a minimum at 220 nm, establishing β-sheet conformation of α-Syn PFFs after sonication. **(E)** Transmission electron microscopy (TEM) of uranyl acetate stained α-Syn PFFs validates the fibrillar ultrastructure of produced fibrils before sonication and reveals a size distribution between 159 and 791 nm, with an average size of 371 nm. **(F)** TEM analysis validates the fibrillar ultrastructure of sonicated fibrils and reveals a size distribution between 37 and 256 nm, with an average size of 123 nm.
